# Incorporating space in hierarchical capture mark recapture models: can we better capture variance?

**DOI:** 10.1101/2022.11.01.514665

**Authors:** Anne – Merel Van Der Drift, Herwig Leirs, Joachim Mariën, Christopher Sabuni, Loth Mulungu, Lucinda Kirkpatrick

## Abstract

Capture mark recapture (CMR) models allow the estimation of various components of animal populations, such as survival and recapture probabilities. In recent years, incorporating the spatial distribution of the devices used to detect an animals’ presence has become possible. By incorporating spatial information, we explicitly acknowledge the fact that there will be spatial structuring in the ecological processes which give rise to the capture data. Individual detection probability is not heterogeneous for a range of different reasons, for example the location of traps within an individual’s home range, the environmental context around the trap or the individual characteristics of an animal such as its age. Spatial capture recapture models incorporate this heterogeneity by including the spatial coordinates of traps, data which is often already collected in standard CMR approaches. Here, we compared how the inclusion of spatial data changed estimations of survival, detection probability, and to some extent the probability of seroconversion to a common arenavirus, using the multimammate mouse as our model system. We used a Bayesian framework to develop non spatial, partially spatial and fully spatial models alongside multievent CMR models. First, we used simulations to test whether certain parameters were sensitive to starting parameters, and whether models were able to return the expected values. Then we applied the non-spatial, partially spatial and fully spatial models to a real dataset. We found that bias and precision were similar for the three different model types, with simulations always returning estimates within the 95% credible intervals. When applying our models to the real data set, we found that the non-spatial model predicted a lower survival of individuals exposed to Morogoro virus (MORV) compared to unexposed individuals, yet in the spatial model survival between exposed and non-exposed individuals was the same. This suggests that the non-spatial model underestimated the survival of seropositive individuals, most likely due to an age effect. We suggest that spatial coordinates of traps should always be recorded when carrying out CMR and spatially explicit methods of analysis should be used whenever possible, particularly as incorporating spatial variation may more easily capture ecological processes without the need for additional data collection that can be challenging to acquire with wild animals.

## INTRODUCTION

When attempting to monitor or manage wild animal populations, particularly elusive or difficult to detect animals, accurate estimations of their abundance and population dynamics are essential (Fuller et al. 2016). Often, this is estimated through capture-mark-recapture (CMR) modelling whereby data obtained from live trapping or other methods of recording the presence of known animals is used to estimate population densities (Kéry and Schaub 2012). A key principle of CMR models is the estimation of the detection probability, which provides the link between the observations of animal presence and true population parameter such as the survival and recapture probability. In CMR studies, animals captured during the initial encounter are marked for identification and released back to the population, after which they are considered available for recapture. Marking makes it possible to collect individual specific information at multiple capture moment whereby numerous capture moments are required to achieve reliable parameter estimations (Chao and Huggins 2010).

Until recently most CMR models excluded the spatial distribution of traps (Royle and Chandler 2014). However, capture mark recapture studies are inherently spatial: traps are laid out in space, and therefore there is spatial structuring in the ecological processes that give rise to the capture data (Royle and Chandler 2014). This is likely to invalidate the assumption of traditional CMR modelling that detection probabilities are assumed to be equal over all individuals (Kéry and Schaub 2012). In practice, the detection probability is heterogeneous due to the spatial organization of individuals relative to traps: the detection probability depends on the number of traps present inside the home range of an individual (Royle and Chandler 2014) and will vary in space as a result of both individual space use and the configuration of detection devices (Dupont et al. 2019a). Spatial capture recapture (SCR) models allow for this individual heterogeneity in detection probabilities (Ergon and Gardner 2014), overcoming the limitations of classical CMR methods without the requirement of additional data, by making use of the spatial trap coordinates that have always been available but rarely utilized. Each individual is associated with a location in space, its activity centre, and the further the activity centre from a detector, the less likely it is that the animal will be detected by that detector (Glennie and Foster 2019). This activity centre is unobserved and therefore considered a latent variable that is estimated by the model, improving the accuracy of the detection model (Glennie and Foster 2019). This improved accuracy in the detection model allows for more accurate estimation of parameters such as density and survival, and therefore makes SCR modelling attractive to use over classical CR modelling (Royle and Chandler 2014). For example Spatially Explicit Capture Recapture (SECR) methods i) find density estimates of 20 – 200% lower than classical CR methods (Obbard, Howe, and Kyle 2010), ii) are more resistant in giving accurate estimates to poor data (Jůnek et al. 2015) and iii) demonstrate that sex differences in survival may be due to spatial processes (Schaub and Royle 2014).

SCR models are often implemented within a Bayesian framework due to its flexibility and accessibility for building complex models, which can be more challenging in frequentist inference (Milleret et al. 2019; Gelman and Hill 2007; Kellner 2018; O’Hagan 2004; Wagenmakers et al. 2008). Additionally, when working with CMR data, traditional frequentist methods can be sensitive to i) low probabilities of detection and survival, ii) small sample sizes and iii) the number of sampling intervals (Calvert et al. 2009). However, it should be recognised that the increased flexibility comes at the expense of being highly computationally demanding (Glennie and Foster 2019), and, due to the inclusion of prior knowledge, can be criticised as being inherently biased. The Bayesian framework allows the user to incorporate the distribution of the data and the distribution of prior knowledge in a natural way (Mariucci, Ray, and Szab 2017), meaning that the posterior distribution is always the result of the chosen prior distribution and the information contained in the data (Kéry and Schaub 2012). The ability to incorporate prior knowledge can result in improved posterior distributions, but if prior knowledge is not truthful, posterior distributions can be deceitful (Kéry and Royle 2016). Therefore, prior selection must be carried out with care, and sensitivity analyses can be used to ensure that the prior does not overwhelm the true information contained within the data. Additionally, diffuse priors can be used which assume no prior knowledge and allow all the information to come from the data.

Due to their increased precision and flexibility, SCR models are increasingly used for estimating population densities of large mammals such as tigers (Dorazio and Karanth 2017), bears (Sun et al. 2017; Gardner et al. 2010) and jaguars (Glennie and Foster 2019), detected by camera trapping (Royle and Chandler 2014) and DNA sampling (Fuller et al. 2016; Royle and Chandler 2014), respectively. Recently, SCR models are also being used to estimate population processes such as within group dispersal and survival through the use of spatial multi-event capture recapture models (Weiser et al. 2018). Multi event capture recapture models not only estimate population densities but also allow for the estimation of transitions between states; for example survival or seroconversion (Sapsford et al. 2015; Hirschinger et al. 2021), allowing users to better understand the role of various aspects of behaviour in state transmission dynamics. This may provide insights into the mechanisms underpinning key processes such as survival and disease transmission. For example, spatial structure plays an important role in plague transmission as *Yersinia pestes* outbreaks are more likely to be initiated by a susceptible animal encountering an infectious carcass than by the transfer of infected fleas (Russell et al. 2021).

Despite the important role that rodents play in ecosystem functioning (Dickman 1999), and the impact they can have on both food production (e.g. agricultural pest species;(Singleton et al. 2015; 2010) and public health (Meerburg, Singleton, and Kijlstra 2009, e.g. as hosts of a range of different zoonotic diseases; plague: Mccauley et al. 2015; Russell et al. 2021, Lassa virus : Frame et al. 1970, and hantaviruses : Guo et al. 2013), there are few examples where spatially explicit capture mark recapture models have been used to investigate their population parameters such as survival (but see e.g. Dupont et al 2021, Casula et al, Romairone et al 2018). Here, we use the multimammate mouse (*Mastomys natalensis*) and Morogoro virus (MORV) as a model system to look at disease transmission. The multimammate mouse, widely distributed through sub-Saharan Africa, causes substantial agricultural damage (H Leirs 1994) and carries a number of zoonotic diseases of public health interest (Weir 2005; WHO 2016). The population dynamics of *Mastomys natalensis* are heavily dependent on rainfall, which is bimodal in this region with long (March-May) and short (November-December) rains. The variation in rainfall results in strong density fluctuations between seasons, generally ranging from 20-300 individuals per hectare (Leirs H. 1994; Sluydts et al. 2009). MORV seems to be endemic in populations of this rodent and can persist even at very low densities, most likely due to the presence of chronically infected animals (Mariën, Borremans, Gryseels, Broecke, et al. 2017; Hoffmann et al. 2021). Based on a previous survival analyses using the same CMR dataset that we will use here, the authors found that MORV-antibody presence was negatively correlated with survival probability and positively with recapture probability of *M. natalensis*. In these models, several confounding factors including, age, season, transience and trap dependence were corrected for. However, spatial factors were not corrected for, despite spatial data being collected. Therefore, this gives an opportunity to explore whether accounting for spatial effects yields similar results to models that also include factors such as age which can be harder to define in wild animals.

The main objective of this study was to explore how incorporating space influences population parameters such as survival probability and transmission dynamics using a combination of real and simulated data. First, we used simulated data to assess how effective our models were at returning the true values, then we ran the models on our field data set to ascertain the extent to which including the spatial information altered our estimates of survival and detection for seropositive and seronegative individuals. This was then compared to the results from Mariën et al (2020) to ascertain the extent to which including spatial variables can capture similar patterns as age inclusive models. Spatial information is explicitly collected in many small mammal CMR studies, yet often not incorporated. However, doing so can often improve density estimates compared to non-spatially explicit approaches (Fauteux et al. 2018). However, to our knowledge this is the first time that the impact of including spatial variables into capture mark recapture models has been used to estimate the impact on state transition dynamics such as infection.

## METHODS

### Study site and data collection

Population and infection data was collected following a standard capture mark recapture method on an open population of *M. natalensis* in Morogoro, Tanzania. A regular grid of 300 traps with traps placed every 10 metres was used. Animals were captured monthly between January 2010 and December 2012 for three consecutive nights with Sherman LFA live traps baited with a mixture of corn flour and peanut butter. Individuals were identified using toe clipping and the sex, mass and reproductive status of each animal was recorded, along with the location of the trap. Blood was sampled to determine the serostate of individuals by using an immunofluorescence assay (IFA) following (Borremans et al. 2015).

The CMR dataset consisted of 6267 total captures of 2512 unique individuals. The individual serostate was used as an indication for current or past infection with Morogoro virus. Based on the results of the IFA, 1314 captured individuals (20.97%) were seropositive. Individuals for which IgG antibodies were not detected in their blood sample were grouped as seronegative individuals and individuals that did show an IgG antibody response were grouped as seropositive individuals. Animals who were first seronegative and seroconverted during the study (0,05% of unique individuals) were also grouped as seropositive individuals, because of the insensitivity of the IFA: antibodies can only be detected 7 days post infection (Borremans et al. 2015). So, when an individual is captured when actually being infected, it can be recorded as seronegative resulting in a false negative result.

#### 2.1 Bayesian statistical modelling (spatial and non-spatial)

To evaluate the CMR data, we created non-spatial and spatial hierarchical models within a Bayesian framework to estimate survival and detection probabilities (ρ and ϕ) of both seropositive and seronegative individuals. Models were fitted in JAGS (Plummer 2016) using R (R Core Team 2018) and the R package jagsUI (Kellner 2018). Both models were applied to simulated and field data. For all analysis, we considered models to have reached convergence based on the 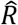 value (Gelman-Rubin test statistic(Gelman and Rubin 1992) < 1.1 and good mixing properties of the different chains which can be assessed by visual inspection of trace and density plots (Gelman and Hill 2007; Kéry and Schaub 2012).

##### 2.1.1 General simulation set-up and procedure

To determine whether models return parameter estimates with acceptable deviation from the truth, we conducted simulations with known true values. To evaluate the performance of the models, we determined the accuracy of the model using measures of bias and precision. Bias was calculated as the difference between the mean of the parameter estimate 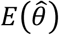 and the set parameter *θ* which was used to simulate the data (Kéry and Schaub 2012). Precision was defined as the standard deviation of the estimated parameter (Kéry and Schaub 2012). Bias and precision were calculated by averaging all the posterior parameter estimations over multiple simulations of each model. All simulations were run with the same following Markov Chain Monte Carlo (MCMC) settings: Three MCMC chains were run for 5000 iterations with a burn-in period of 500 iterations and a thinning rate of 10.

##### 2.1.2. Non-spatial CR model simulation set-up

Simulated data for the non-spatial CR model was created using the function `simul.ms` provided in (Kéry and Schaub 2012). In the simulation dataset 160 individuals were present and 800 individuals were captured in total whereby individuals can be in three states (seropositive, seronegative and dead), similar to the real dataset. We simulated 6 monthly capture sessions where captures took place for 3 consecutive days. The survival probability (ϕ_A_) was set to 0.6 for seronegative individuals and to 0.5 for seropositive individuals (ϕ_B_).The transition from seronegative to seropositive (ψ_AB_) was set to 0.2 and the transition from seropositive to seronegative (ψ_BA_) was constrained to be 0.005, as in our system transition from positive to negative is considered very unlikely and reflects insensitivity in the immunofluorescence assay used to detect antibodies against MORV rather than waning immunity (Borremans et al. 2015). In addition, the detection probability of seronegative individuals (ρ_A_) was set to 0.5 while for seropositive individuals the detection probability (ρ_B_) was set to 0.6. Data was simulated 1000 times to evaluate the performance of the model and average estimates and standard errors across all simulations were compared to the true value. Bias and precision were calculated for all of the parameters above except for detection. In addition, a parameter sensitivity test was carried out to analyse how sensitive the non-spatial non temporal model is to different parameter values (0.2, 0.4, 0.6, or 0.8). This test was carried out by varying the parameters of survival (ϕ) and detection (ρ) for both seronegative and seropositive individuals and the transition probability from seronegative to seropositive (ψ_BA_).

##### 2.1.3 Spatial CR model simulation set up

Simulated data for the spatial CR model was created by the simulation function `simul.sCJSN` provided in (Schaub and Royle 2014). Originally 100 individuals were present in the population at *t* = 0 and 800 individuals were captured in total in a study grid of 15 by 15. Animals were captured at 7 capture moments. The survival probability (ϕ) was set to be 0.6, the detection probability (ρ) was set to be 0.5 and the dispersal variances in both X and Y direction (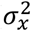 and 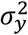) were set to be 0.5. Data was simulated 1000 times to evaluate the performance of the model by conducting an accuracy test without adding the group variable to make a distinction between seropositives and seronegatives. Bias and precision are calculated for all parameters: ϕ, ρ and 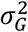.

#### 2.2 Non-spatial CR model

For the purpose of the first analysis, we created a multistate non-spatial model to allow for the estimation of survival (ϕ), detection (ρ) and transition from seronegative to seropositive (ψ_AB_). As previous experiments have confirmed that lifelong immunity follows infection with MORV (Borremans et al. 2015), the transition from seropositive to seronegative (ψ_BA_) was constrained to be minimal by using a beta distributed prior with *α* = 1 and *β* = 50. This causes more than 99% of the prior distribution to fall between 0 and 0.02, rendering the probability of transitioning from seropositive to seronegative to effectively zero. If we fully constrain ψ_BA_ to be 0, the model gives up too much flexibility and fails to converge. Additionally, we cannot discount the possibility that a minority of animals may sero-revert over time. All priors of the other parameters were allocated a uniform non-informative distribution.

The multistate non-dispersal model consisted of two sub-models:

(a) **The state model**, which describes the state of an individual at time t+1, given its state at time t. The state model can be described by a set of state equations:

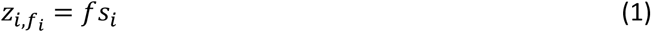

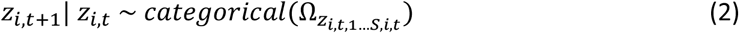

In equation 1, the true state of individual *i* is described by element 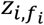, which indicates the true state of individual *i* at its first encounter. We assume that an individual can only be observed in their actual state (absence of state assignment error), therefore, *f_Si_* is identical to the observed state at first encounter 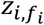 (Kéry and Schaub 2012).

We recognise three true states: alive and seronegative (state A), alive and seropositive (state B) and dead. Theoretically, 9 different transitions can occur, but due to the restriction of transition ψ_BA_, this is unlikely. We define all possible transitions in a four-dimensional state-transition matrix (Ω, eq. 2). The first dimension describes the state at time *t*, the second dimension the state at *t* + 1, the third dimension the individual (*i*) and the fourth dimension describes time (*t*). We use a categorical likelihood to account for the multistate character of the model (Kéry and Schaub 2012).

(b) **The observation model**, which links the true state to the observed state. This model is described by the following equation:

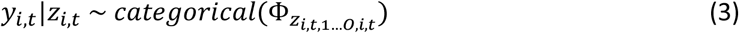

Where the observed state *y* is conditional on the true state *z*. Similar to the true state transition matrix, an observation matrix (Θ) is created. This matrix is also four-dimensional whereby the first dimension states the true state, the second the observed state, the third the individual *i* and the fourth dimension describes time *t*. Similar to the number of true states, the number of observed states also equals three: seen as seronegative, seen as seropositive and not seen.

#### 2.3 : Partial dispersal model

We compared a partial dispersal model (in that rodents were grouped by dispersal distances) to assess whether accounting for movement by simple grouping was sufficient to allow for similar estimates of survival as that from the fully spatial model. In this model, we were also able to estimate transition probabilities between states. Therefore, we compared parameter estimates of survival (ϕ), detection (ρ) and transition (ψ_AB_) between short and long dispersing animals. Individuals were grouped based on the maximal dispersal distance with a cut-off of 40 metres (equ. 4). The same parameters as in the first analysis are of interest: ϕ_A_, ϕ_B_, ψ_AB_, ρ_A_ and ρ_B_.

Since capture locations are recorded, it is possible to calculate dispersal distances based on X and Y coordinates when individuals are captured at least two times. If an individual is captured more than twice, the two traps that are the farthest apart are chosen to calculate the maximal individual dispersal distance, calculated as the Euclidean distance.

Animals with a maximal dispersal distance shorter than 40 metres were grouped as short dispersing animals, whereas animals with a dispersal distance longer than 40m were grouped as long dispersing animals. The cut-off was chosen based on data from previous CMR and radio-tracking studies (Borremans et al. 2014; Herwig Leirs, Verheyen, and Verhagen 1996). In CMR studies, mean day-to-day movement of 20 meters are observed whereby 85% of the individuals do not move more than 25 meters (Borremans et al. 2014; Herwig Leirs, Verheyen, and Verhagen 1996). However, in radio tracking studies, larger day-to-day dispersal distances are observed. This is due to short excursions that cannot be captured in CMR studies possibly due to i) interception by traps before the excursion or ii) trap shyness during the excursion (Herwig Leirs, Verheyen, and Verhagen 1996). Based on previous CMR and radio tracking studies and to account for individual variation, we therefore decided on a cut-off between small and long dispersing animals at 40 metres, although we recognise that this is somewhat arbitrary. With a cut-off of 40 metres, 65% of the individuals in the field dataset were grouped as short dispersing animals and 35% as long dispersing animals.

We used the same model structure as the multistate non dispersal model (2.2), with the addition of a grouping variable. We used model 1 without any fixed or random time effects for parameter estimates as individual grouping was based on dispersal distances over all capture moments, so it was therefore it is not justifiable to make parameter estimates of survival, detection or transition with time as a fixed or random effect.

The MCMC settings for both the simulated and the field data set were the same as above (2.2). To evaluate the performance of the multistate partial dispersal model, bias and precision were calculated twice for the parameters ϕ_A_, ϕ_B_, ψ_AB_, ψ_BA_, ρ_A_ and ρ_B_ for both groups of short and long dispersing animals.

#### 2.4 Spatial CR model

Similar to the non-spatial CR model, the spatial Cormack – Jolly – Seber (sCJS) CR model also consists of two sub-models:

a) **The state model**, which describes the state of an individual at time t+1, given its state at time t. The state model can be described by two different state processes. The first state process, described by equations 4 and 5, indicates whether an individual is alive (*z_i,t_* = 1) or dead (*z_i,t_* = 0). Individuals are grouped as seronegative or seropositive whereby individuals that were ever seropositive are considered as always seropositive. State transitions are not estimated in the spatial model due to parameter redundancy, however survival and dispersal are estimated separately for seropositive and seronegative individuals (Weiser et al. 2018). As in the non-spatial CR model, the latent state variable is conditional on first capture which is indicated by *f_i_*. All priors in this spatial model are set to be non-informative and therefore a uniform distribution is assigned as a prior for all parameters.

Element *s_i,t_* in eq. 5 shows the true survival probability between *t* and *t* + 1 for individual *i*. We assume that survival is the same inside and outside the study area with no spatial variation. We also assume that *z_i,t_* is individually independent and is conditional on the survival probability *s_i,t_* (Schaub and Royle 2014). The likelihood of this model is based on a Bernoulli distribution instead of a categorical distribution in the non-spatial model, because the model only contains two states: alive and dead.

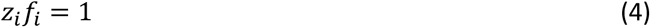

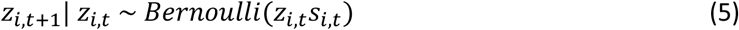

Equation 6 describes the second state process pertaining to the dispersal location variable *G*. As individual dispersing behaviour can change between *t* and *t* + 1 the location at *t* + 1 is modelled with the normal distribution with i) estimated mean *G_i,t_*, which indicates the location of individual *i* at time *t* and ii) a variance 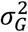 which indicates the dispersal variance in X and Y direction (Schaub and Royle 2014).

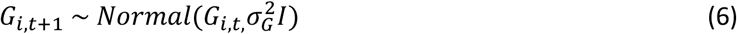

(b) **The observational model**, which links the two state processes with the observed state. In the multistate models, observation *y* was only conditional on the true state data *z_i,t_*. However, since in this model dispersal is included which is modelled with a normal distribution, animals are also able to disperse outside the study area. Individuals are only available for recapture when being alive and present in the study area, and therefore the observation is conditional on both. *r_i,t_* indicates if the location *G_i,t_* of individual *i*, who is alive, is inside or outside the study area *A* at time *t*. *r_i,t_* = 0, when a living animal is outside the study area at time *t* and *r_i,t_* = 1, when a living animal is inside the study area at time *t* (eq. 7 and 8, (Weiser et al. 2018). Dispersal distances are not modelled directly, but dispersal variances in X and Y direction will be estimated. Based on the estimated variances, the dispersal distance can be calculated (eq. 4).

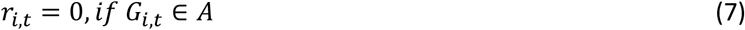

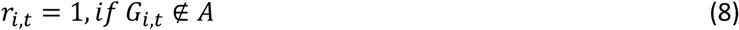

Knowing this, the observation model can be described by equation 9. The likelihood of the observational model is a Bernoulli distribution with the parameters *z_i,t_*, *r_i,t_* and *p_i,t_* whereby *p_i,t_* is the detection probability of individual *i* at time *t*. Because this model is parameter redundant, it is assumed that survival and recapture probabilities are similar among individuals and are constant over time (Schaub and Royle 2014).

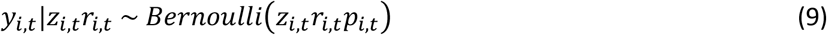

We provide example JAGS code (Supplementary material) that was developed from published examples (referenced within example JAGS code). For the non spatial and partial dispersal models used a burn-in period of 5000 iterations and a thinning rate of 10 which consistently provided good mixing across three chains. We ran both models for 75,000 iterations, resulting in 7500 saved iterations which were then used to generate posterior distributions of parameters. For the full dispersal model, we used the same burn-in and thinning rates, but only needed to run the model for 25,000 iterations to achieve convergence, as excluding state transition matrices reduces the variance in the model.

## RESULTS

### 3.1 Survival and transition probabilities from the non-spatial CJS model

Initially we investigated whether allowing for time variance in either survival, detection or transmission probability resulted in a better model fit, using the non-spatial CJS model (table 1). Two models failed to converge and were excluded from further analyses. The best fitting model assumed a constant probability of survival and transition from seronegative to seropositive but a fixed time effect on detection probability (table 1, model 2; Figure 1). We found that the survival probability of seronegative individuals (0.562 with CRI_95%_= [0.544, 0.580]) was higher compared to survival of seropositive individuals (0.485 with CRI_95%_= [0.450, 0.520], Figure 1A). The overlap between both survival distributions is 0.46%, suggesting that seropositivity is associated with a significantly lower survival probability. We estimated the transition probability from seronegative to seropositive to be 0.077 (CRI_95%_= 0.065, 0.091, figure 1B). Monthly detection probabilities estimated for seronegative and seropositive individuals are shown in figure 1C, demonstrating no difference in detection probability over time between the two groups.

**Table 1:**
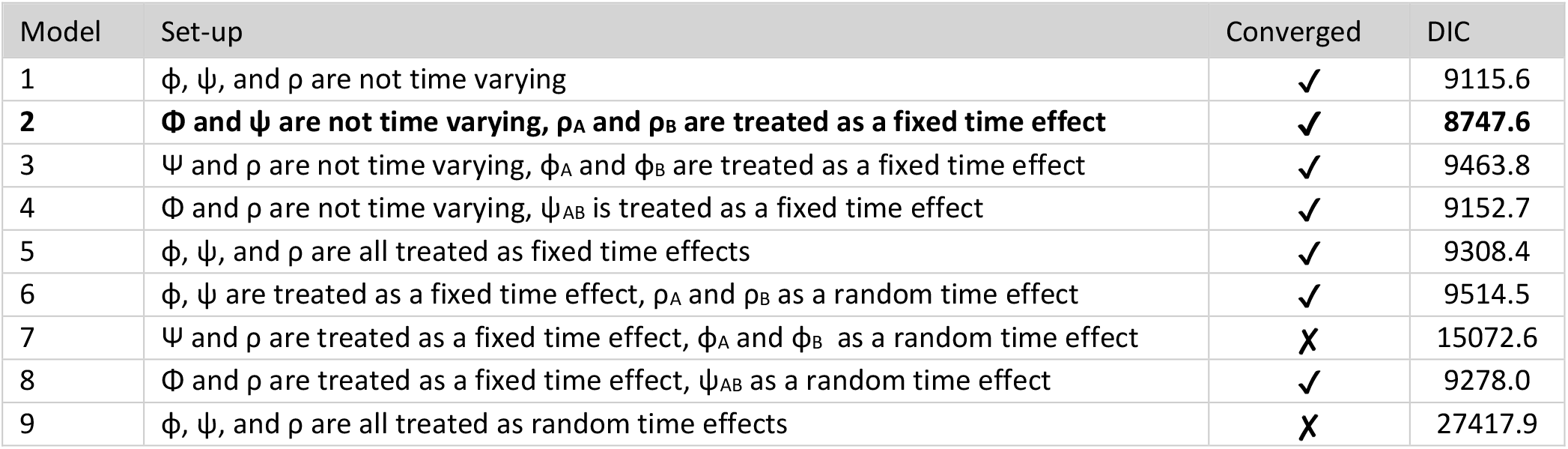
Comparison of non-spatial CR models where survival, detection and transition from seronegative to seropositive were allowed to vary with no, fixed or random time effects.

| Model | Set-up | Converged | DIC |
| --- | --- | --- | --- |
| 1 | $\phi$, $\psi$, and $\rho$ are not time varying | ✓ | 9115.6 |
| 2 | **$\Phi$ and $\psi$ are not time varying, $\rho_A$ and $\rho_B$ are treated as a fixed time effect** | ✓ | **8747.6** |
| 3 | $\Psi$ and $\rho$ are not time varying, $\phi_A$ and $\phi_B$ are treated as a fixed time effect | ✓ | 9463.8 |
| 4 | $\Phi$ and $\rho$ are not time varying, $\psi_{AB}$ is treated as a fixed time effect | ✓ | 9152.7 |
| 5 | $\phi$, $\psi$, and $\rho$ are all treated as fixed time effects | ✓ | 9308.4 |
| 6 | $\phi$, $\psi$ are treated as a fixed time effect, $\rho_A$ and $\rho_B$ as a random time effect | ✓ | 9514.5 |
| 7 | $\Psi$ and $\rho$ are treated as a fixed time effect, $\phi_A$ and $\phi_B$ as a random time effect | ✗ | 15072.6 |
| 8 | $\Phi$ and $\rho$ are treated as a fixed time effect, $\psi_{AB}$ as a random time effect | ✓ | 9278.0 |
| 9 | $\phi$, $\psi$, and $\rho$ are all treated as random time effects | ✗ | 27417.9 |

**Figure 1:**
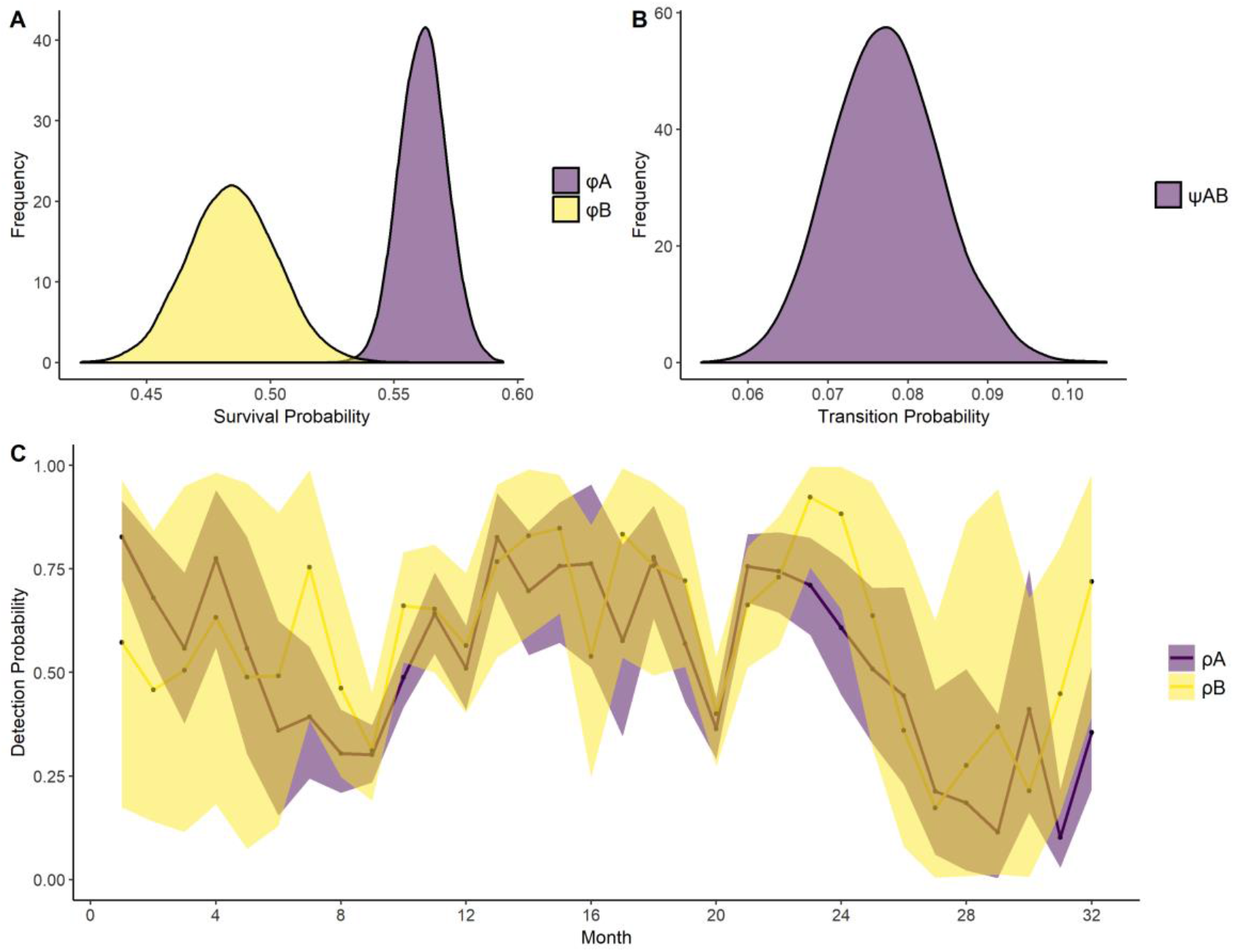
Posterior distributions of estimated population parameters. (A) The full distribution of the estimated survival probabilities for seronegative individuals (φA shown in purple) and seropositive individuals (φB shown in yellow). (B) The full distribution of the estimated transition probability from seronegative (state A) to seropositive (state B). (C) The estimated detection probabilities for both seronegative (ρA) and seropositive individuals (ρB) at each monthly capture moment. Estimations are shown by the points and the shaded areas present the associated CRI

In order to assess the accuracy of the model, bias and precision were calculated for survival (ϕ_A_ and ϕ_B_) and the transition probability (ψ_AB_) from seronegative to seropositive (table 2) for the best fitting model using simulated data with known true parameter values. Because of the time-dependency of the detection probability, bias and precision could not be calculated for this parameter. We carried out a parameter sensitivity test to analyse how sensitive the model was to different parameter values (0.2, 0.4, 0.6, or 0.8), using the non-temporal model (model 1, table 1) in order to be able to compare with the partial spatial model (Figure 2). The precision of parameter estimates was greatest when survival and detection probabilities were high and when transition probabilities (ψ_AB_) were low (table 2). However, in all cases the true value is within the credible intervals returned by the model on the simulated data, giving reassurance that the model performs well and returns reliable estimates for the non-spatial CR model (figure 2).

**Table 2:** *Bias and precision calculated from the simulated models for survival* (ϕ*), detection (*ρ*) and dispersal variance (*σ*) for the non spatial, partially spatial (grouped according to maximal movement distance) and fully spatial (not grouped by serostate in the simulation model). No estimates for* ρ *for the non spatial model are available due to the inclusion of a fixed effect for time*.

| Model | Parameter | Bias | Precision |
| --- | --- | --- | --- |
| Non spatial | $\phi_A$ | -0.005 | 0.030 |
| | $\phi_B$ | 0.001 | 0.032 |
| | $\psi_{AB}$ | 0.007 | 0.028 |
| | $\rho_A$ | NA | NA |
| | $\rho_B$ | NA | NA |
| Partially spatial | $\phi_A$ | -0.002 | 0.027 |
| | $\phi_B$ | 0.002 | 0.030 |
| | $\psi_{AB}$ | 0.006 | 0.027 |
| | $\rho_A$ | 0.008 | 0.043 |
| | $\rho_B$ | -0.005 | 0.049 |
| Fully spatial | $\phi$ | -0.001 | 0.022 |
| | $\rho$ | 0.002 | 0.032 |
| | $\sigma$ | 0.210 | 0.029 |

**Figure 2.**
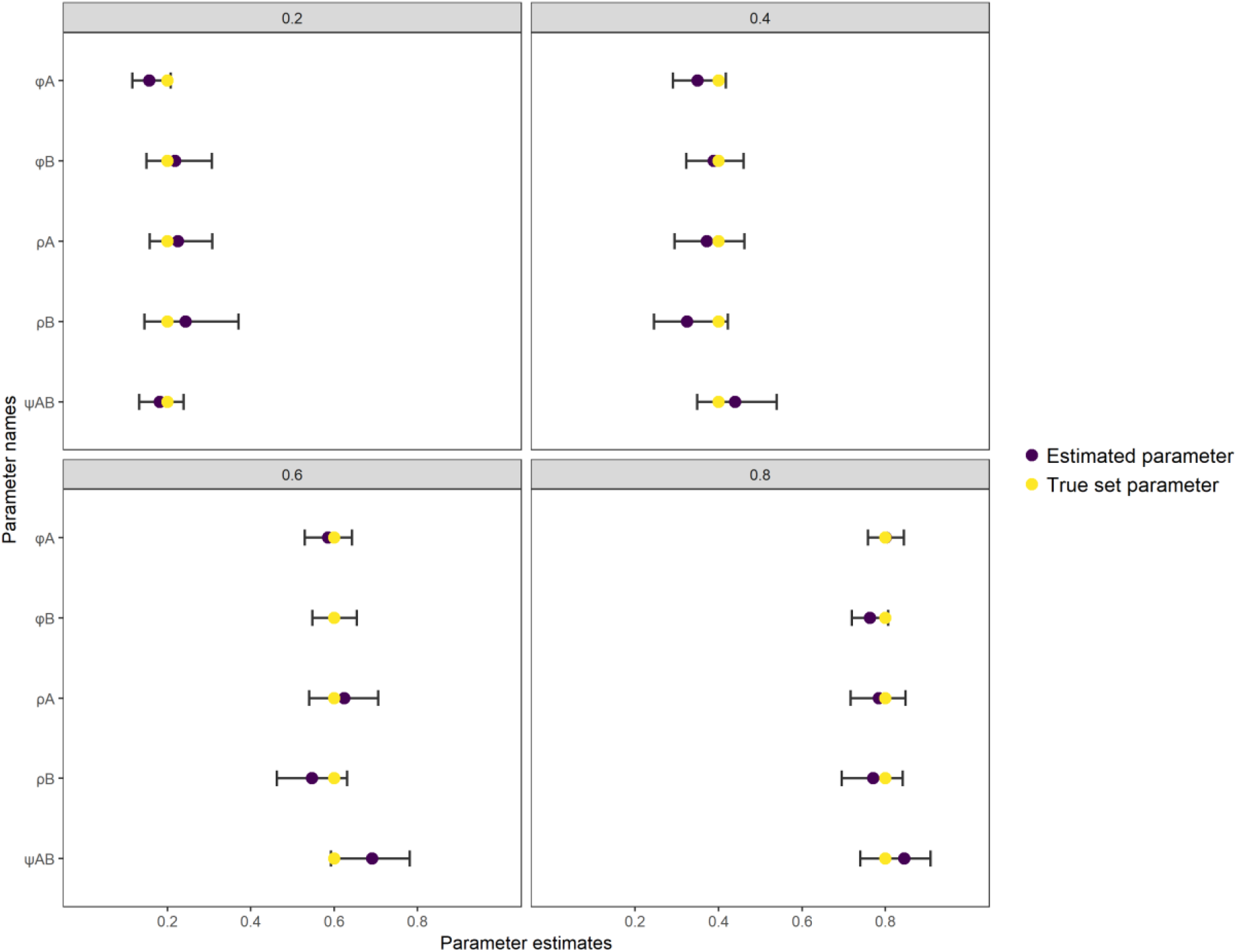
Sensitivity test of all parameters of interest of the non-spatial CR model. Every parameter of interest (transition, survival and detection) was set to be 0.2, 0.4, 0.6 or 0.8 in the simulated dataset which are corresponding to the four different graphs. Parameter estimations are made for every simulated data set whereby the true set parameter (yellow) can be compared to the estimated parameter (purple)

#### Inclusion of movement

We included movement parameters in two different ways; first we created a grouped model where we looked at survival, detection and transition probability for animals that were grouped by either short or long distance movements. Next, we used a fully spatial model where we assessed survival, detection and movement for seropositive and seronegative individuals. Due to models becoming over parametrised we were unable to assess transition probabilities in a spatial multistate model, hence our use of the two different model types. The partial movement model gave estimates with the highest accuracy for survival and the lowest accuracy for detection regardless of the serostate, as shown in table 2. Similar to the results for the non-spatial model, we found no evidence that the model was sensitive to different initial true parameters; in all simulations the true value fell within the estimated 95% BCI. The fully spatial model gave estimates with the highest accuracy for detection and the lowest accuracy for survival, although bias was similarly low for both estimates. It should be taken into account that survival and detection are estimates of probabilities and dispersal is an estimate of variance. Therefore, the accuracy of dispersal cannot be compared to the accuracy of the other parameters.

When movement was included in the partial dispersal model, we found that the transition probability for short dispersing animals (0.077 with CRI_95%_= [0.062, 0.094) is comparable with the transition probability for long dispersing animals (0.081 with CRI_95%_= [0.058, 0.107]; Fig 3), with an overlap between transition probability distributions of 64.5%. Similar to the non-spatial model, survival of seropositive individuals is lower than for seronegative individuals, particularly for long distance moving animals (Fig 3 B). For seropositive individuals, survival of individuals that moved less than 40m (0.628 with CRI_95%_= [0.585, 0.670]) is higher than individuals that moved more than 40m 0.561 with CRI_95%_= [0.486, 0.635], Fig 3B).

**Figure 3:**
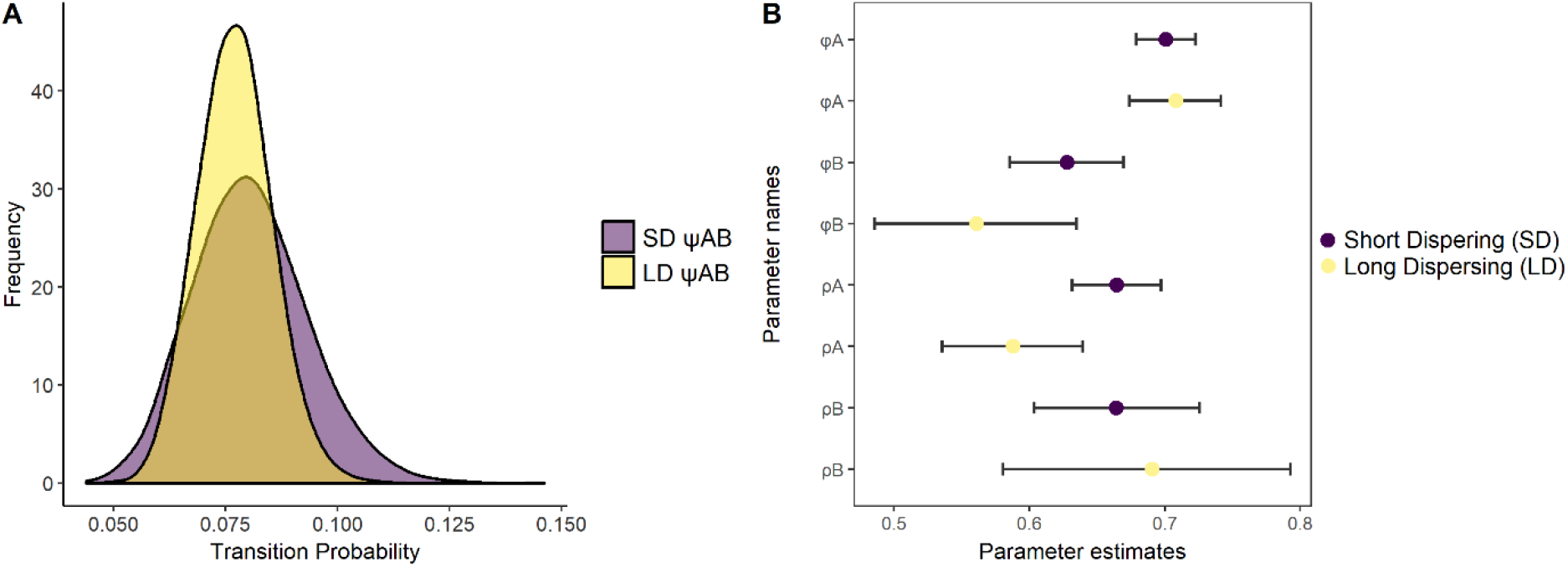
Posterior distributions of estimated population parameters. (A) The estimated transition probability from seronegative (state A) to seropositive (state B) for short and long dispersing animals (shown in purple and yellow). (B) Parameter estimates are shown for survival of seronegatives (ϕ_A_) and seropositives (ϕ_A_) and detection of seronegatives (ρB) and seronegatives (ρB) for both short and long dispersing animals (SD and LD). Estimated points are shown by the points and the lines are representing the 95% credible intervals.

In contrast to the non-spatial and partially spatial CR models, the spatial CR model found no difference in survival of seronegative and seropositive individuals (Mean seronegative: 0.594, CRI_95%_= [0.575, 0.613] and mean seropositive: 0.584, CRI_95%_= [0.546, 0.621] respectively, Figure 3B, 4B). Similarly, detection probability for seronegative individuals was similar to that of seropositive individuals (Mean seronegative: 0.636 with CRI_95%_= [0.602, 0.670], mean seropositive: 0.675 with CRI_95%_= [0.610 0.738], Figure 3B). The dispersal variance for seronegative individuals 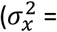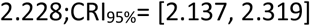 and 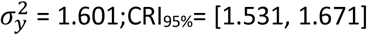, Figure 3A) tended to be higher than the variance for seropositive individuals (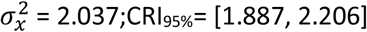 and 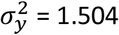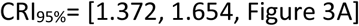, Figure 3A]). The estimated dispersal distance for seronegative individuals is 21.90 meters with CRI_95%_= [20.98, 22.81] compared to a distance of 20.20 meters with CRI_95%_= [18.612, 21.999] for seropositive individuals. In comparison to the non-spatial and partially-spatial models, fully incorporating movement into the model design fundamentally altered estimates of survival, with there no longer being any difference in survival based on serostate (Figure 5)

**Figure 4:**
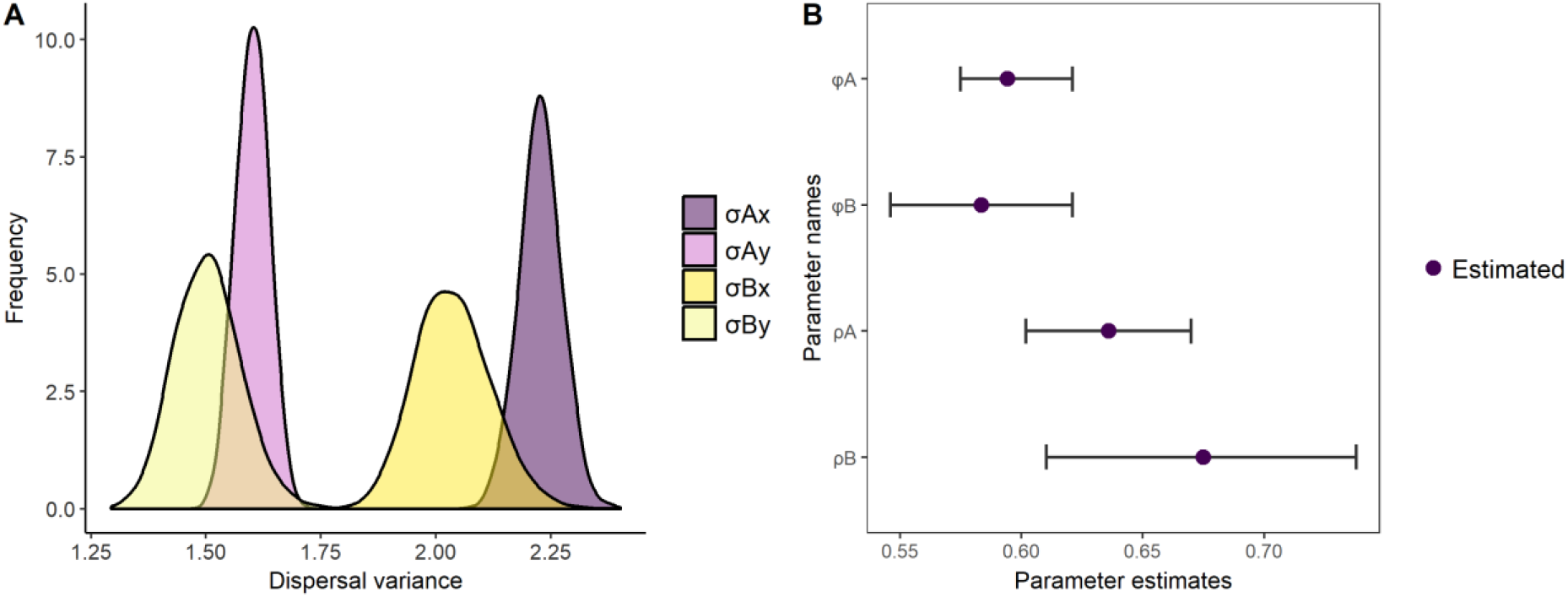
Posterior distributions of estimated population parameters from the full dispersal model. (A) The estimated dispersal variances in X and Y direction for both seronegative and seropositive individuals. (B) Parameter estimates are shown for survival and detection of seronegatives (a) and seronegatives (b). Estimates are shown by the purple points and the lines are representing the 95% credible intervals.

**Figure 5:**
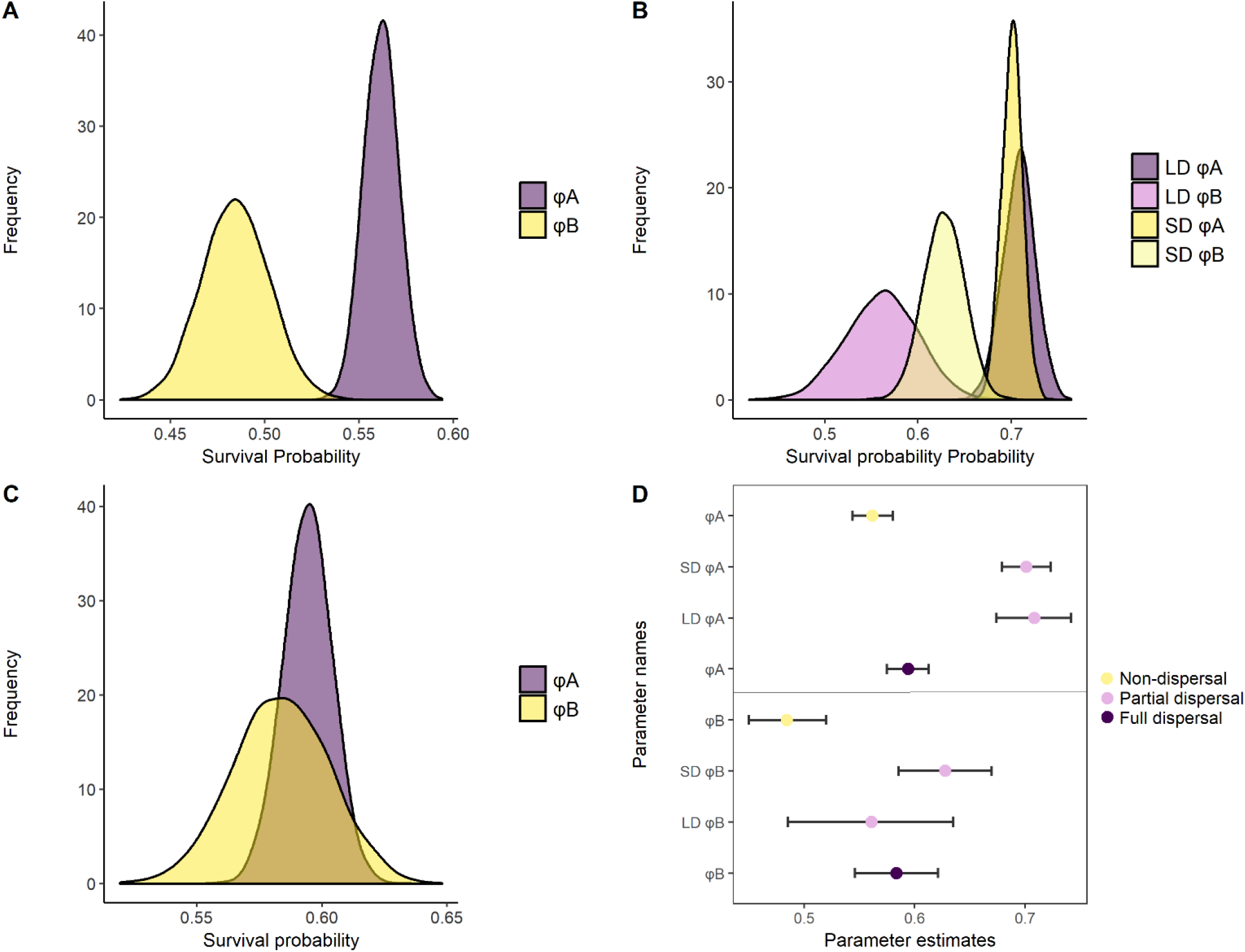
Survival probabilities estimated by different models are compared with each other. (A) Survival probabilities for seronegative (ϕ_A_) and seropositive (ϕ_B_) individuals estimated by the multistate non-spatial model. (B) Survival probabilities for seronegative and seropositive individuals grouped as short (SD) or long (LD) distance movement animals estimated by the multistate partial spatial model. (C) Survival probabilities for seronegative (ϕ_A_) and seropositive (ϕ_B_) individuals estimated by the full dispersal model. (D) All estimated survival probabilities of the three different models plotted together in one graph.

Comparing the non-spatial and spatial model estimates for survival probability for seronegative and seropositive individuals showed that when spatial movements are not included in the model, survival in seropositive individuals appeared to be far lower than in seronegative individuals. However, after including spatial effects, this difference was greatly reduced (Figure 5A-C).

## DISCUSSION

Capture mark recapture methods have had a profound influence on our understanding of demography and population processes. However, many of these advances have occurred independent of and unconnected to the spatial structure of the either the population or the landscape within which the population is situated (Royle, Fuller, and Sutherland 2018). Therefore, CMR methods have historically been non spatial, accounting for neither the inherent spatial nature of sampling or the spatial distribution of individual encounters (Royle, Fuller, and Sutherland 2018). The advent of spatially explicit capture mark recapture models (Chandler and Andrew Royle 2013; Jůnek et al. 2015; Royle and Chandler 2014; Ergon and Gardner 2014) represents an important advancement in our ability to quantify and explicitly study the role of spatial processes. Here, we demonstrate that including spatial information in standard CR models can have a significant effect on population parameter estimates such as survival. Therefore, when spatial data exists, as is the case with the vast majority of capture mark recapture, attempts should be made to include this in model construction.

From our sensitivity analysis, we found that most of the estimated survival probabilities are underestimated in the non-spatial model. This is also reflected in the field data analysis: when using traditional non-spatial CR modelling, survival estimates are lower compared to survival estimates from inclusion of spatial movement information. It is likely that parameter estimates are underestimated due to non-recorded movement outside of the study area (Schaub and Royle 2014) and the fact that capture probabilities are unequal among all traps, despite classical non-spatial CR models assuming the opposite (Kéry and Schaub 2012). Due to the individual specific location of home ranges, some traps will occur in more home ranges than other traps, therefore making the capture probability variable across traps (Royle and Chandler 2014). In contrast, when accounting for space in spatial CR models, equal detection probabilities are not assumed, rather this variation in detection probabilities is now modelled. Spatial capture recapture models assume that animal activity centres are uniformly and independently distributed over the state space and that the area therefore includes all the animals available for sampling (López-Bao et al. 2018). It is likely that this assumption is violated to some degree in our system, as environmental conditions vary across the study grid and intraspecific interactions may also play a role (Bischof et al. 2020). However, due to a lack of territoriality in *M. natalensis* (Borremans et al. 2014), and evidence that SCR models appear to be fairly robust to violations of this assumption (López-Bao et al. 2018), it is unlikely that this has significant bearing on our results. Finally, SCR models are sensitive to sparse data, which can lead to imprecise estimates. Therefore, SCR models are more appropriate for systems where multiple detections of the same individual occurs at different locations, in order to estimate the location of latent activity centres (Dupont et al. 2019).

Comparing between the classical non-spatial CR model and the spatial CR model, survival of seropositive individuals (ϕB) is underestimated more than survival of seronegative individuals (ϕA). This might be explained by the difference in group size between seronegative (n=1996) and seropositive (n=516) individuals, whereby less information is contained within the data for seropositive compared to seronegative individuals. Alternatively, these differences could be due to an age effect. Firstly, we record seroprevalence rather than active infection, therefore older individuals are more likely to be positive simply because they have had more time to become exposed (Borremans et al. 2011). Secondly, dispersal behaviour may differ between age groups. Juvenile *M. natalensis* are more exploratory than adults (Vanden Broecke et al. 2017), and therefore may disperse further in order to establish their own home range away from the natal area and related conspecifics (Van Hooft et al. 2008). In the study of Mariën et al. (2018), age was corrected for by using the logarithm of body weight on the first capture, and by only retaining individuals who weighed less than 35g at their first capture session as these were considered to be juveniles. A seasonal effect of survival was found, alongside a limited impact of exposure status (Mariën 2018). In our study we included animals that were presumed adult at first capture, with the assumption that age and seasonal effects are accounted for by movement patterns (Vanden Broecke et al. 2017). Therefore, we cannot separate whether the differences we see are due to either seasonal or age related differences. Given that infection with MORV does not appear to impact *M. natalensis* body condition (Mariën, Borremans, Gryseels, Soropogui, et al. 2017), an impact of prior exposure to MORV on survival seems unlikely (Mariën et al. 2018). If, as hypothesised in Mariën et al (2017), exposure to MORV is associated with movement behaviour in the mice (e.g bolder mice are more likely to become infected and trapped; Vanden Broecke et al. 2017, or infection with MORV influences individual behaviour and trap happiness; Vyas 2015), this would be captured in the spatial CMR model and may also explain why we no longer see a difference in survival estimates given prior exposure to MORV. In addition, detection probabilities and dispersal differences were similar for both seronegative and seropositive individuals in the fully spatial model, suggesting that the impacts of trap affinity are captured in the spatial model and that exposure to MORV does not impact trap affinity and movement.

An alternative explanation would be that the difference in underestimation between groups is due to differential distribution of home ranges between the two groups, and that exposure to MORV is related to unknown spatial processes. For example, the home ranges of the seropositive individuals might be located closer to toward the edges of the study grid suggesting a relatively lower detection probability compared to the seronegative individuals. This lower detection probability instructs mortality rather than emigration. This theory is supported by the so-called edge effect, which states that animals will more frequently be captured in the traps close to the edge compared to the traps located in the centre a study grid (Stenseth and Hansson 1979). This edge effect also applies to the studied *M. natalensis* population in Morogoro in Tanzania (H Leirs 1994). In this study area, it is observed that at the edges at certain capture occasions, the traps are more occupied than in the centre, possibly as a consequence of occasional trapped individuals that are passing by the grid. The incidence of these passers-by is larger at the edges compared to the centre of the grid giving the appearance of a higher density at the edge of the grid (H Leirs 1994). When animals are more frequently captured at the edges of the study grid, this indirectly indicates more undetected movement leading to a higher underestimation of survival. Analysing the influence of this edge-effect on population parameter estimations might be of interest for future research. However, there is little evidence to suggest that spatial variation at the edge of the grid compared to the centre of the grid would result in differential probabilities of exposure to MORV, especially as transmission is considered to be predominantly from direct contact or from within nest encounters (e.g mating or as an outcome of breeding attempts from chronically infected animals, Hoffmann et al. 2021). In addition, SCR models only require that the distribution of detectors is described by the encounter model, rather than adhering to any particular layout, and appear to be fairly robust to mis-specification of the encounter probability model (Fuller et al. 2016).

We found no difference in transition probability when comparing between the non-spatial and partly spatial models, and no impact of movement on the probability of seroconversion. We were not able to include state transitions in the fully spatial models but these results suggest that there is little impact of spatial processes on seroconversion, and that seroconversion itself may be a hyperlocal event. Given the small dispersal distances predicted from our model, which were in line from other studies (Borremans et al. 2014; Herwig Leirs, Verheyen, and Verhagen 1996), this suggests that most exposure risk is likely to occur in the immediate area of occupancy of the individual.

In this study we did not correct for the number of trapping events per individual through the use of a robust design. This may influence the estimated dispersal distance, since animals that are trapped more frequently have a higher probability of having a longer dispersal distance. However, even without the use of a robust design model, we find similar results from our simple non spatial CR model compared to the frequentist models of Mariën et al. (2018), where age, seasonal effects and trap dependence are accounted for within a robust design. The results of the spatial model suggest that accounting fully for movement may be more effective than accounting for other effects such as age; movement may therefore be the way in which these grouping parameters are influencing animal behaviour. Spatial data may also be easier to include than individual factors such as age which can be hard to estimate in wild animal populations.

Here, we demonstrate that accounting for dispersal and movement is key when estimating population parameters in a capture mark recapture study and can have a dramatic impact on parameter estimates. While recent advances in maximum likelihood approaches to estimate density offer considerable improvements over Bayesian approaches in terms of processing time (Glennie and Foster 2019), Bayesian approaches provide a more flexible and easily interpretable framework within which to construct these models. We suggest that where possible, inclusion of spatial parameters should be preferred when using CR models to investigate survival and transmission dynamics in a population, and that the kind of capture mark recapture studies that are often used for rodent research lend themselves well to this kind of approach (Romairone et al. 2018). Future approaches that allow for the characterisation of state transition dynamics while also accounting for space would be a valuable addition when investigating processes such as disease transmission.

