## Supplementary for "Incorporating space in hierarchical capture mark recapture models: can we better capture variance?"

**NON DISPERSAL MODEL**

Jags model R code for the general non-dispersal model used for the first and second analysis described in 2.5 and 2.6. The functions used to simulate capture mark recapture data are also included in the code. Code taken from Schaub and Royle 2012

```
# Setup simulation parameters
phiA <- 0.6
phiB <- 0.5
psiAB <- 0.2
psiBA <- 0.05
pA <- 0.5
pB <- 0.6
n.occasions <- 6                                          #Number of occasions
n.states <- 3                                             #Number of states
n.obs <- 3                                                #Number of observations
marked <- matrix(NA, ncol = n.states, nrow = n.occasions) #Number of released     individuals
marked[,1] <- rep(100, n.occasions)
marked[,2] <- rep(60, n.occasions)
marked[,3] <- rep(0, n.occasions)

# Generation of two four-dimensional matrices
# 1. Generation of the state-transition matrix
totrel <- sum(marked)*(n.occasions-1)
PSI.STATE <- array(NA, dim=c(n.states, n.states, totrel, n.occasions-1))
for (i in 1:totrel){
  for (t in 1:(n.occasions-1)){
    PSI.STATE[,,i,t] <- matrix(c(
      phiA*(1-psiAB), phiA*psiAB,     1-phiA,
      phiB*psiBA,     phiB*(1-psiBA), 1-phiB,
      0,              0,              1       ), nrow = n.states, byrow = TRUE)
  } #t
} #i

# 2.Generation of the Observation matrix
PSI.OBS <- array(NA, dim=c(n.states, n.obs, totrel, n.occasions-1))
for (i in 1:totrel){
  for (t in 1:(n.occasions-1)){
    PSI.OBS[,,i,t] <- matrix(c(
      pA, 0,  1-pA,
      0,  pB, 1-pB,
      0,  0,  1       ), nrow = n.states, byrow = TRUE)
  } #t
} #i

# Define function to simulate multistate capture-recapture data
simul.ms <- function(PSI.STATE, PSI.OBS, marked, unobservable = NA){
  # Unobservable: number of state that is unobservable
  n.occasions <- dim(PSI.STATE)[4] + 1
  CH <- CH.TRUE <- matrix(NA, ncol = n.occasions, nrow = sum(marked))
  # Define a vector with the occasion of marking
  mark.occ <- matrix(0, ncol = dim(PSI.STATE)[1], nrow = sum(marked))
  g <- colSums(marked)
  for (s in 1:dim(PSI.STATE)[1]){
    if (g[s]==0) next  # To avoid error message if nothing to replace
    mark.occ[(cumsum(g[1:s])-g[s]+1)[s]:cumsum(g[1:s])[s],s] <-
      rep(1:n.occasions, marked[1:n.occasions,s])
  } #s
  for (i in 1:sum(marked)){
    for (s in 1:dim(PSI.STATE)[1]){
      if (mark.occ[i,s]==0) next
      first <- mark.occ[i,s]
      CH[i,first] <- s
      CH.TRUE[i,first] <- s
    } #s
    for (t in (first+1):n.occasions){
      # Multinomial trials for state transitions
      if (first==n.occasions) next
      state <- which(rmultinom(1, 1, PSI.STATE[CH.TRUE[i,t-1],,i,t-1])==1)
      CH.TRUE[i,t] <- state
      # Multinomial trials for observation process
      event <- which(rmultinom(1, 1, PSI.OBS[CH.TRUE[i,t],,i,t-1])==1)
      CH[i,t] <- event
    } #t
  } #i
  # Replace the NA and the highest state number (dead) in the file by 0
  CH[is.na(CH)] <- 0
  CH[CH==dim(PSI.STATE)[1]] <- 0
  CH[CH==unobservable] <- 0
  id <- numeric(0)
  for (i in 1:dim(CH)[1]){
    z <- min(which(CH[i,]!=0))
    ifelse(z==dim(CH)[2], id <- c(id,i), id <- c(id))
  }
  return(list(CH=CH[-id,], CH.TRUE=CH.TRUE[-id,]))
  # CH: capture histories to be used
  # CH.TRUE: capture histories with perfect observation
}

# Define the number of simulations and data structure to store output
nsim <- 100
phiA.est <- phiB.est <- psiAB.est <- psiBA.est <- pA.est <- pB.est <- numeric()

# Start loop for the simulation
for (sim in 1:nsim){

# Execute function and store capture history (CH)
sim_sim <- simul.ms(PSI.STATE, PSI.OBS, marked)
CH <- sim_sim$CH

# Compute vector with occasion of first capture
get.first <- function(x) min(which(x!=0))
f <- apply(CH, 1, get.first)

# Recode CH matrix
# 1 = seen alive in A, 2 = seen alive in B, 3 = not seen
rCH <- CH
rCH[rCH==0] <- 3

# Function to create known latent states z
known.state.ms <- function(ms, notseen){
  # notseen: label for 'not seen'
  state <- ms
  state[state==notseen] <- NA
  for (i in 1:dim(ms)[1]){
    m <- min(which(!is.na(state[i,])))
    state[i,m] <- NA
  }
  return(state)
}

# Function to create initial values for unknown z
ms.init.z <- function(ch, f){
  for (i in 1:dim(ch)[1]){ch[i,1:f[i]] <- NA}
  states <- max(ch, na.rm = TRUE)
  known.states <- 1:(states-1)
  v <- which(ch==states)
  ch[-v] <- NA
  ch[v] <- sample(known.states, length(v), replace = TRUE)
  return(ch)
}

# Model 1: No time variation, but including restricted psiBA
sink("model1.jags")
cat("
    model {
    ####################################################
    # Parameters:
    # phiA: survival probability at state A
    # phiB: survival probability at state B
    # psiAB: movement probability from state A to state B
    # psiBA: movement probability from state B to state A
    # pA: recapture probability at state A
    # pB: recapture probability at state B
    ###################################################
    # States (S) as defined in state-transition matrix
    # 1 alive in state A
    # 2 alive in state B
    # 3 dead
    # Observations (O) as defined in observational matrix
    # 1 seen in state A
    # 2 seen in state B
    # 3 not seen
    ###################################################
    # State A is defined as being seronegative
    # State B is defined as being seropositive
    ####################################################
    # Priors and constraints
    for (t in 1:(n.occasions-1)){
    phiA[t] <- mean.phi[1]
    phiB[t] <- mean.phi[2]
    psiAB[t] <- mean.psiAB
    psiBA[t] <- mean.psiBA
    pA[t] <- mean.p[1]
    pB[t] <- mean.p[2]
    }

    mean.psiAB ~ dunif(0,1)   # Non-informative prior
    mean.psiBA ~ dbeta(1,50)  # psiBA is restricted based on prior knowledge

    for (u in 1:2){
    mean.phi[u] ~ dunif(0, 1) # Non-informative prior
    mean.p[u] ~ dunif(0, 1)   # Non-informative prior

    # Define parameters
    for (i in 1:nind){

    # Define probabilities of state S(t+1) given S(t)
    for (t in f[i]:(n.occasions-1)){ #loop over time
    # First index = states at time t-1, Second index = individual specific,
    # Third index = time specific, Last index = states at time t
    ps[1,i,t,1] <- phiA[t] * (1-psiAB[t])
    ps[1,i,t,2] <- phiA[t] * psiAB[t]
    ps[1,i,t,3] <- 1-phiA[t]
    ps[2,i,t,1] <- phiB[t] * psiBA[t]
    ps[2,i,t,2] <- phiB[t] * (1-psiBA[t])
    ps[2,i,t,3] <- 1-phiB[t]
    ps[3,i,t,1] <- 0
    ps[3,i,t,2] <- 0
    ps[3,i,t,3] <- 1

    # Define probabilities of O(t) given S(t)
    # First index = states at time t, Second index = individual specific,
    # Third index = time specific, Last index = observations at time t
    po[1,i,t,1] <- pA[t]
    po[1,i,t,2] <- 0
    po[1,i,t,3] <- 1-pA[t]
    po[2,i,t,1] <- 0
    po[2,i,t,2] <- pB[t]
    po[2,i,t,3] <- 1-pB[t]
    po[3,i,t,1] <- 0
    po[3,i,t,2] <- 0
    po[3,i,t,3] <- 1
    } #t
    } #i

    # State-space model likelihood
    for (i in 1:nind){
    z[i,f[i]] <- y[i,f[i]]
    for (t in (f[i]+1):n.occasions){ #loop over time
    # State process: draw S(t) given S(t-1)
    z[i,t] ~ dcat(ps[z[i,t-1], i, t-1,])
    # Observation process: draw O(t) given S(t)
    y[i,t] ~ dcat(po[z[i,t], i, t-1,])
    } #t
    } # i
    }
    ",fill = TRUE)
sink()

# Bundle data
bugs.data <- list(y = rCH, f = f, n.occasions = dim(rCH)[2], nind = dim(rCH)[1], z = known.state.ms(rCH, 3))

# Initial values for all parameters of interest
inits.mod1 <- function(){list(mean.phi = runif(2, 0, 1), mean.psiAB = runif(1, 0, 1), mean.psiBA = rbeta(1, 1, 50), mean.p = runif(2, 0, 1), z = ms.init.z(rCH, f))}
# Parameters monitored
parameters.mod1 <- c("mean.phi", "mean.psiAB", "mean.psiBA", "mean.p")

# MCMC settings
ni <- 75000   #Number of iterations
nt <- 10      #Thinning rate
nb <- 5000    #Burn-in period
nc <- 3       #Number of chains

#call JAGS
mod1_it <- jags(bugs.data, inits.mod1, parameters.mod1, "model1.jags", n.chains = nc, n.thin = nt, n.iter = ni, n.burnin = nb, parallel = TRUE, n.cores = 3)

# Store simulation results
phiA.est[sim] <- mod1_b_p$mean$mean.phiA
phiB.est[sim] <- mod1_b_p$mean$mean.phiB
psiAB.est[sim] <- mod1_b_p$mean$mean.psiAB
psiBA.est[sim] <- mod1_b_p$mean$mean.psiBA
pA.est[sim] <- mod1_b_p$mean$mean.pA
pB.est[sim] <- mod1_b_p$mean$mean.pB
} #sim
```

**FULL DISPERSAL MODEL**

Jags model code for the full dispersal model described in 2.7. The functions used to simulate capture mark recapture data are also included.

```
# Input parameters:
# rel: number of newly released individuals at each occasion
# x.space: dimension of study area in x-direction (x-limits are 0 and x-space)
# y.space: dimension of study area in y-direction (y-limits are 0 and y-space)
# nyears: number of capture occasions
# s: survival probability (we assume here that it is constant over time)
# p: recapture probability (we assume here that it is constant over time)
# disp.varX: variance of the dispersal distribution in x-direction
# disp.varY: variance of the dispersal distribution in y-direction

# Output parameters:
# Y.in: capture-recapture data sampled within the study area
# Y.comp: all capture-recapture data (i.e. not restricted to the study area)
# G.in: X and Y-coordinates of the encountered individuals within the study area
# G.comp: X and Y-coordinates of all encountered individuals (i.e. not restricted
# to the study area)
# f: vector with the release occasion for each individual
simul.sCJSN <- function(rel = 20, x.space = 15, y.space = 15, nyears = 20, s = 0.5,
                        p = 0.6, disp.varX = 5, disp.varY = 5){
  n <- rel * (nyears-1) # total number of individuals
  Z <- d <- matrix(NA, ncol = nyears, nrow = n)
  G <- G.in <- G.comp <- G.in.comp <- array(NA, dim=c(n, nyears, 2))
  f <- rep(1:(nyears-1), rep(rel, (nyears-1))) # vector indicating when
  #individuals were captured for the first time
  # Simulating survival process
  for (i in 1:n){
    Z[i,f[i]] <- 1
    for (t in (f[i]+1):nyears){
      Z[i,t] <- rbinom(1, 1, s) * Z[i,t-1]
    } # t
  } # i
  # Simulating dispersal process
  # determine site of first capture
  for (i in 1:n){
    G[i,f[i],1] <- runif(1, 0, x.space)
    G[i,f[i],2] <- runif(1, 0, y.space)
  } # i
  # dispersal (assuming a normal distribution in both directions)
  for (i in 1:n){
    for (t in (f[i]+1):nyears){
      G[i,t,1] <- G[i,t-1,1] + rnorm(1, 0, sqrt(disp.varX))
      G[i,t,2] <- G[i,t-1,2] + rnorm(1, 0, sqrt(disp.varY))
    } # t
  } # i
  # only keep coordinates when an individual was alive
  G[,,1] <- G[,,1] * Z
  G[,,2] <- G[,,2] * Z
  G[G==0] <- NA

  # Simulating sampling
  # Capture histories
  Y <- Z
  for (i in 1:n){
    for (t in (f[i]+1):nyears){
      Y[i,t] <- rbinom(1, 1, p) * Z[i,t]
    } # t
  } # i
  # Exclude locations that are outside the boundary (study area)
  inside <- Z
  excl <- which(G[,,1] < 0 | G[,,1] > x.space | G[,,2] < 0 | G[,,2] > y.space)
  inside[excl] <- 0
  # Capture-recapture data within study area
  Y.in <- Y * inside
  G.in[,,1] <- G[,,1] * inside * Y
  G.in[,,2] <- G[,,2] * inside * Y
  G.in[G.in==0] <- NA
  # Capture-recapture data including all captures (not restricted to study area)
  Y.comp <- Y
  G.comp[,,1] <- G[,,1] * Y
  G.comp[,,2] <- G[,,2] * Y
  G.comp[G.comp==0] <- NA
  # Output
  return(list(Y.in = Y.in, Y.comp = Y.comp, G.in = G.in, G.comp = G.comp, f = f))
}

# Function that is required for creating initial values (from Kery & Schaub 2012)
known.state.cjs <- function(ch){
  state <- ch
  for (i in 1:dim(ch)[1]){
    n1 <- min(which(ch[i,]==1))
    n2 <- max(which(ch[i,]==1))
    state[i,n1:n2] <- 1
    state[i,n1] <- NA
  }
  state[state==0] <- NA
  return(state)
}

# Full dispersal model
sink("sCJSN.mod")
cat("
    model {
    # Priors and constraints
    for (i in 1:nind){
    for (t in f[i]:(n.occasions-1)){
    phi[i,t] <- mean.phi
    p[i,t] <- mean.p
    } #t
    } #i
    mean.phi ~ dunif(0,1)
    mean.p ~ dunif(0,1)
    for (i in 1:2){
    tau[i] <- pow(sigma[i], -2)
    sigma[i] ~ dunif(0, 50)
    }
    # Likelihood
    for (i in 1:nind){
    # Define latent state at first capture
    z[i,f[i]] <- 1
    for (t in (f[i]+1):n.occasions){
    # State processes
    # Survival
    z[i,t] ~ dbern(phi[i,t-1] * z[i,t-1])
    # Dispersal
    G[i,t,1] ~ dnorm(G[i,t-1,1], tau[1])
    G[i,t,2] ~ dnorm(G[i,t-1,2], tau[2])
    # Observation process
    # Test whether the actual location is in- or outside the state-space
    r[i,t] <- step(G[i,t,1]-Xmin) * step(Xmax-G[i,t,1]) * step(G[i,t,2]-Ymin) *
    step(Ymax-G[i,t,2])
    Y[i,t] ~ dbern(p[i,t-1] * z[i,t] * r[i,t])
    } #t
    } #i
    }
    ",fill = TRUE)
sink()

# Define the number of simulations and data structure to store output
nsim <- 1000
phi.est <- p.est <- sigma.est <- DIC.est <- numeric()

# Start loop for the simulation
for (sim in 1:nsim){

dat <- simul.sCJSN(rel = 100, x.space = 15, y.space = 15, nyears = 7, s = 0.6, p =
                       0.5, disp.varX = 0.5, disp.varY = 0.5)

# Bundle data
jags.data <- list(Y = dat$Y.in, G = dat$G.in, Xmin = 0, Xmax = 15, Ymin = 0, Ymax =
                      15, f = dat$f, nind = nrow(dat$Y.in), n.occasions = ncol(dat$Y.in))

# Initial values for all parameters of interest
inits <- function(){list(mean.phi = runif(1,0,1), mean.p = runif(1,0,1), sigma =
                             runif(2,0.01,2), z = known.state.cjs(dat$Y.in))}
# Parameters monitored
parameters <- c("mean.phi", "mean.p", "sigma")

# MCMC settings
ni <- 5000    #Number of iterations
nt <- 10      #Thinning rate
nb <- 500     #Burn-in period
nc <- 3       #Number of chains

#Call JAGS
library(R2jags)
schaub_simulated_1000 <- jags(jags.data, inits, parameters, "sCJSN.mod", n.chains = nc, n.thin = nt,
                                n.iter = ni, n.burnin = nb)
# Store results
phi.est[sim] <- schaub_simulated_1000$mean$mean.phi
p.est[sim] <- schaub_simulated_1000$mean$mean.p
sigma.est[sim] <- schaub_simulated_1000$mean$sigma
DIC.est[sim] <- schaub_simulated_1000$DIC
} #sim
```
